# Soft sweeps and beyond: Understanding the patterns and probabilities of selection footprints under rapid adaptation

**DOI:** 10.1101/114587

**Authors:** Joachim Hermisson, Pleuni S Pennings

## Abstract

1. The tempo and mode of adaptive evolution determine how natural selection shapes patterns of genetic diversity in DNA polymorphism data. While slow mutation-limited adaptation leads to classical footprints of “hard” selective sweeps, these patterns are different when adaptation responds quickly to a novel selection pressure, acting either on standing genetic variation or on recurrent new mutation. In the past decade, corresponding footprints of “soft” selective sweeps have been described both in theoretical models and in empirical data.
2. Here, we summarize the key theoretical concepts and contrast model predictions with observed patterns in *Drosophila,* humans, and microbes.
3. Evidence in all cases shows that “soft” patterns of rapid adaptation are frequent. However, theory and data also point to a role of complex adaptive histories in rapid evolution.
4. While existing theory allows for important implications on the tempo and mode of the adaptive process, complex footprints observed in data are, as yet, insufficiently covered by models. They call for in-depth empirical study and further model development.

## 1 Hard and soft selective sweeps

The view of adaptation in molecular evolution has long focused almost exclusively on a mutation-limited scenario. The assumption in such a scenario is that beneficial mutations are rare, so that they are unlikely to be present in the population as standing genetic variation (SGV) or to occur multiple times in a short window of time. Mutation-limited adaptation therefore occurs from single new beneficial mutations that enter the population only after the onset of the selection pressure (e.g. due to environmental change). When such a beneficial mutation fixes in the population, it reduces the genetic diversity at linked neutral loci according to the classical model of a “hard” selective sweep (Maynard Smith and Haigh, 1974; Kaplan et al., 1989; Barton, 1998). If recurrent beneficial mutation is considered at all in a mutation-limited scenario, each such mutation is assumed to create a new allele. Single adaptive steps then either proceed independently of each other or compete due to clonal interference, where adaptation is slowed by linkage (Gerrish and Lenski, 1998; Desai and Fisher, 2007). Despite of evidence for rapid adaptation from quantitative genetics and phenotypic studies (Messer et al., 2016), SGV or recurrent origins of the same allele have long been ignored in molecular evolution.

Around a decade ago, several publications started to explore selective sweeps outside the mutation-limitated scenario (Innan and Kim, 2004; Hermisson and Pennings, 2005; Przeworski et al., 2005; Pennings and Hermisson, 2006a,b). These papers described novel patterns for genetic footprints termed “soft sweeps”. It has since become clear that non-mutation-limited adaptation and soft sweeps are probably much more common than originally thought. However, recent years have also seen some lively debate about whether soft sweeps are everywhere (Messer and Petrov, 2013) or rather a chimera (Jensen, 2014). Some aspects of this dispute root in diverging interpretation of the available data, others go back to conceptual differences. With this review, we aim to provide an overview and intuitive understanding of the relevant theory and the patterns that are observed in model species.

## Definitions

Selective sweeps refer to patterns in genomic diversity that are caused by recent adaptation. Characteristic patterns (footprints) arise if rapid changes in the frequency of a beneficial allele, driven by positive selection, distort the genealogical history of samples from the region around the selected locus. We can therefore understand sweep footprints via properties of their underlying genealogy or *coalescent* history (Wakeley, 2008). In this context, the key genealogical implication of mutation-limited adaptation is that, at the selected locus itself, the time to the most recent common ancestor (MRCA) of the sample, *T*_MRCA_, is shorter than the time that has elapsed since the onset of the new selection pressure, *T_S_*. For recent adaptation, we thus obtain coalescent histories that are much shorter than the expected neutral coalescent time *T_N_* (2*N_e_* generations for a pair of lineages in a diploid population of effective size *N_e_*). It is this shortened genealogy that is responsible for the characteristic footprint of a hard sweep (see the Footprints section below).

- We define a hard sweep based on the sample genealogy of the beneficial allele, requiring that (i) *T*_MRCA_ ≪ *T_N_* (recent adaptation), *and* (ii) *T*_MRCA_ ≤ T_S_ (a single recent ancestor), see Fig. 1A.

**Figure 1:**
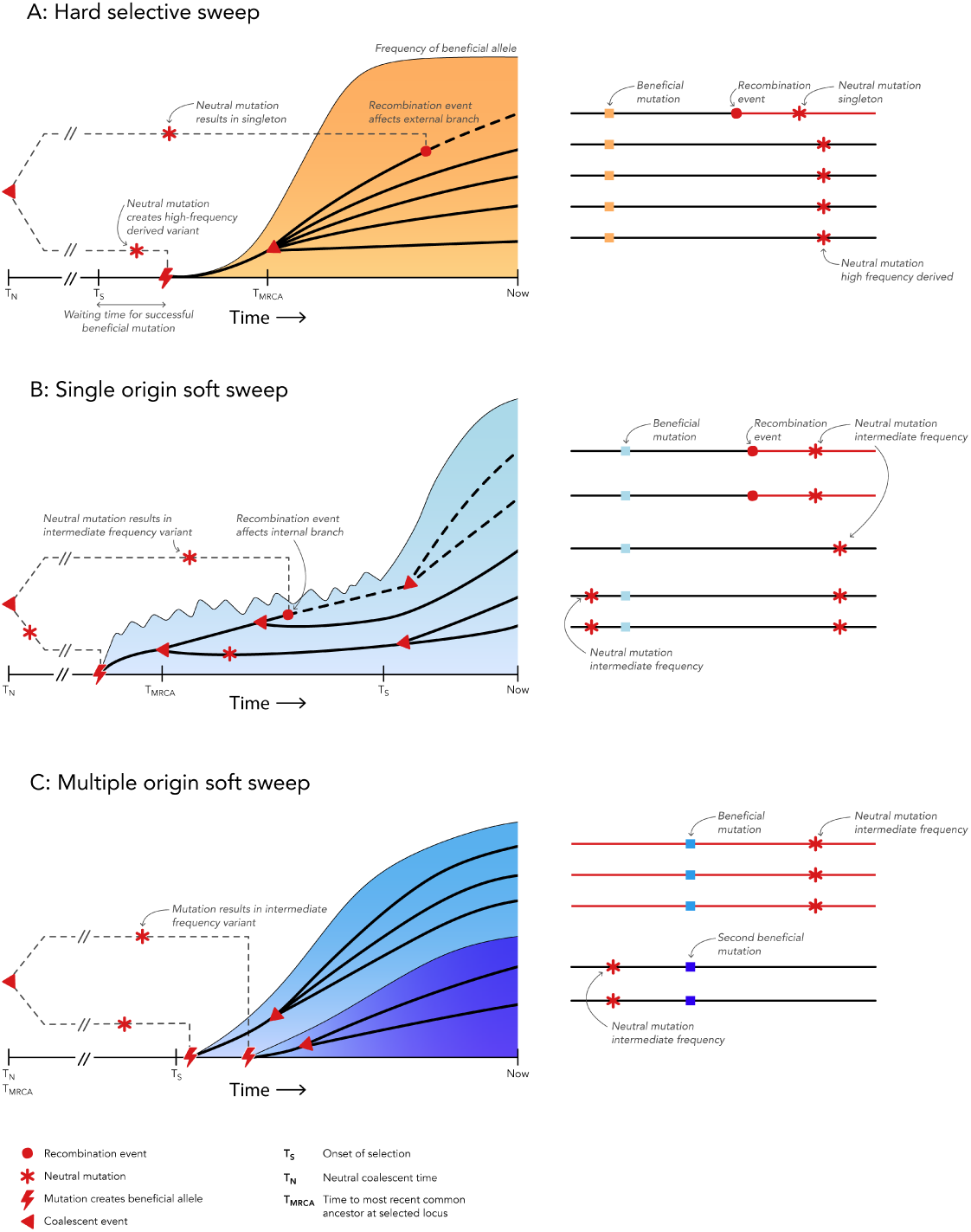
Hard and soft sweep types. Colored regions mark the frequency of those copies of the beneficial allele that still have descendants at the time of sampling. Black and dashed lines show coalescent histories at linked sites. On the right, mutations and recombination events are also shown on haplotypes of the five sampled individuals. (A) For a hard sweep, the time to the most recent common ancestor at the selected site *T*_MRCA_ is shorter than the time since the onset of selection *T_S_*. All ancestral variation at tightly linked sites is eliminated. Recombination leads to low-frequency and high-frequency derived variants in flanking regions. (B) For a single-origin soft sweep from the SGV, multiple lines of descent of the beneficial allele reach into the “standing phase” before *T_S_*. Early recombination introduces ancestral haplotypes at intermediate frequencies. (C) The beneficial allele traces back to multiple origins. Each origin introduces an ancestral haplotype, typically at intermediate frequency.

If adaptation is not mutation limited because the beneficial allele was already present in the population prior to the onset of selection, or because the mutation occurs recurrently, both assumptions of a hard sweep can be violated: The time to the MRCA typically predates the onset of the selection pressure, *T*_MRCA_ > *T_S_*, and can approach the neutral coalescent time, *T*_MRCA_ ~ *T_N_*. Still, if adaptation is recent, we can observe a characteristic footprint: a soft selective sweep.

- For a soft sweep, we require that (i) *T_S_* ≪ *T_N_* (recent adaptation), and (ii) *T*_MRCA_ > *T_S_* (more than a single recent ancestor) for the sample genealogy of the beneficial allele.

There are two different ways how soft sweep genealogies can come about, which lead to different patterns (Hermisson and Pennings, 2005).

1. *Single-origin soft sweeps* refer to a genealogy that traces back to a single mutational origin of the beneficial allele, but comprises multiple copies of this allele in the SGV when positive selection sets in at time *T_S_*, see Fig. 1B.
2. *Multiple-origin soft sweep:* In this case, the sample genealogy of the beneficial allele comprises multiple origins of this same allele from recurrent mutation (or from some other source like migration). These origins can occur prior to the onset of selection (standing variation), but also only after that (new variation), Fig. 1C.

For both, hard and soft sweeps, the definitions apply irrespective of whether the beneficial allele has reached fixation (complete sweep) or still segregates in the population (partial sweep), as long as we restrict the sample to carriers of the beneficial allele only. Note that, for a given sweep locus, we may observe a soft sweep in one sample, but a hard sweep in a different sample. The probability to observe a soft sweep increases with sample size, up to the whole population. We discuss below under which conditions sample size has (or has not) a major influence on this probability.

Two non-exclusive processes can lead to soft sweeps, which both capture essential aspects of non-mutation-limited adaptation: adaptation from SGV and adaptation from recurrent mutation. However, there is no one-to-one correspondence of a process and the type of selective sweep that results, see Fig. (2). In particular, adaptation from SGV can either lead to a hard sweep (if a single copy from the standing variation is the ancestor of all beneficial alleles in the sample), or to a soft sweep, which can either be *single-origin* or *multiple-origin.* The notion of a “soft sweep” as defined here following previous work (e.g. Hermisson and Pennings, 2005; Messer and Petrov, 2013; Jensen, 2014; Berg and Coop, 2015) is therefore not synonymous for “adaptation footprint from SGV”. Sweep types refer to classes of patterns that result from characteristic coalescent genealogies, not to evolutionary processes. This leaves us the task to explore the *probability* of each sweep type under a given evolutionary scenario and to use this information for statistical inference of process from pattern.

## 2 Footprints of hard and soft sweeps

In order to distinguish footprints of hard and soft sweeps, we need to understand how these footprints are shaped by the different characteristic features of the underlying genealogies. We assume the following model. A diploid population of constant size *N_e_* experiences a new selection pressure at time *T_S_*. The derived allele *A* is generated from the wildtype *a* with fitness 1 by recurrent mutation of rate *u.* The frequency of *A* is *x* = *x*(*t*). Prior to *T_S_*, *A* is neutral or deleterious with fitness disadvantage 1 – *s_d_*, *s_d_* ≥ 0. After time *T_S_*, allele A is beneficial with fitness 1 + *s_b_*. For simplicity, we assume codominance.

### Footprints of hard sweeps

The hallmark of a hard sweep genealogy is a very recent common ancestor of all beneficial alleles in the sample. Among carriers of the beneficial allele, ancestral variation prior to the onset of selection can only be preserved if there is recombination between the polymorphic locus and the selection target. Between the closest recombination breakpoints to the left and to the right of the selected allele, we obtain a core region without ancestral variation. In the flanking regions, ancestral variation is maintained on some haplotypes due to recombination. These *recombination haplotypes* generate characteristic signals in the site-frequency spectrum.

We can understand the characteristics of hard sweep footprints from the shape of typical genealogies as sketched in Figure 1A: The genealogy directly at the selected site is “star-like”, with all coalescence events happening in a short time interval when the frequency x of the beneficial allele is very small. This is because all ancestors (between now and the MRCA) in a hard sweep genealogy must carry the beneficial allele. The number of potential ancestors at any given point in time is 2*N_e_x.* For small *x*, the coalescence probability 1/(2*N_e_x*) thus becomes very large (Barton, 1998).

The average fixation time of a beneficial allele with selection coefficient s_b_ is ≈ 2log[4*N_e_sb*]*/sb* generations (van Herwaarden and van der Wal, 2002; Hermisson and Pennings, 2005). All recombination events that can restore ancestral variation in a sample need to occur during this phase. For strong selection, this time is short so that we obtain a broad core region without any variation – other than new mutations that occur during or after the sweep. Its width is inversely proportional to the recombination rate *r* times the fixation time and thus roughly ~ *s_b_*/*r* (Kaplan et al., 1989). Due to the star-like genealogy, recombination during the selective phase will typically only affect single lineages, which (back in time) have not yet coalesced with other lineages (Fig. 1A). Recombination therefore introduces single long branches into the genealogy of a linked neutral locus, which reach far back into the time prior to the selective phase. Neutral mutations that occur on these branches either lead to low-frequency variants or high-frequency derived variants, depending on whether they occur on the internal or external long branch (Fig. 1A). This explains the typical site-frequency spectrum in the flanking regions of hard sweeps with an excess of high- and low-frequency alleles (Braverman et al., 1995; Fay and Wu, 2000). The recombination haplotypes create positive linkage disequilibrium (LD) in the flanking regions of a hard sweep. However, since separate recombination events are needed to both sides, there is (on average) no LD across the selected site (Kim and Nielsen, 2004; Stephan et al., 2006).

### Footprints of single-origin soft sweeps

For single-origin soft sweeps, like for hard sweeps, all beneficial alleles in the sample trace back to a single origin. Coalescent histories at the selected site are therefore confined to the descendants of the founding mutation (the colored region in Fig. 1B). In contrast to hard sweeps, however, the MRCA predates the onset of selection, *T*_MRCA_ > *T_S_* (measured from “Now”). This is possible if adaptation occurs from SGV. If the allele is neutral or only slightly deleterious as it arises, it can stay in the population for a long time, its frequency governed by drift. Only when the allele becomes beneficial, it quickly rises in response to selection.

The pattern of a single-origin soft sweep depends on the age of the MRCA relative to the onset of selection. If *T*_MRCA_ is not much older than *T_S_*, the pattern will look like a hard sweep. This is typically the case if the selected allele is strongly deleterious in the ancestral environment, because deleterious alleles do not stay in the population for a long time. On the other hand, if *T*_MRCA_ is much older than *T_S_*, a distinct footprint of a single-origin soft sweep will be visible. If the fitness of the selected allele is 1 – *s_d_* before *T_S_* and 1 + *s_b_* after *T_S_*, this will be the case for alleles with a weak fitness trade-off, *s_d_* ≪ *s_b_*. Hermisson and Pennings (2005) define a parameter of relative selective advantage,

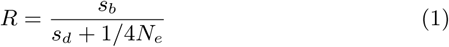

and estimate that a clear pattern of a single-origin soft sweep (relative to a hard sweep with strength *s_b_* throughout) can be expected for R ≳ 100.

Alleles with a weak trade-off (large R) have a larger chance to be picked up from SGV and then will typically lead to soft sweeps (see the Probabilities section below). Patterns have been described in detail for adaptation from neutral variation (*s_d_* = 0) by Przeworski et al. (2005), Peter et al. (2012) and Berg and Coop (2015). The essential difference relative to hard sweeps is that genealogies are no longer star-like because the frequency *x* of the (later) beneficial allele changes only slowly during the “standing phase” prior to *T_S_*. The order of coalescence, recombination, and neutral mutation events during this phase is unaffected by *x*. As a consequence, early mutation and recombination events often affect multiple individuals and lead to intermediate-frequency polymorphism in a sample (Fig. 1B). Berg and Coop (2015) show that the number and frequencies of these early recombination haplotypes follow the Ewens sampling distribution (Ewens, 1972). The net effect is a weakening of the sweep signal, especially close to the selected site (where the hard sweep signal is strongest): the core region is narrower and nearby flanking regions are not strongly dominated by low-frequency variants.

### Footprints of multiple-origin soft sweeps

A soft sweep genealogy with multiple origins of the beneficial allele extends into the part of the population that carries the ancestral allele (Fig. 1C), where coalescence happens on a neutral time scale. This leads to the key characteristic of this sweep type: a pattern of extended haplotypes (one per origin) that stretch across the selected site. In contrast to the recombination haplotypes of a hard sweep, haplotypes corresponding to different mutational origins are typically observed at intermediate frequencies in population samples. Like the early recombination haplotypes of a single-origin soft sweep, they follow a Ewens distribution (Pennings and Hermisson, 2006a).

The difference in expected frequencies of recombination haplotypes of hard sweeps and mutation haplotypes of multiple-origin soft sweeps can be understood as follows. Imagine we follow a lineage back in time. In each generation, it “picks” an ancestor from the 2*N_e_x* beneficial alleles in that generation. The probability that this ancestor is a new beneficial mutant is 2*N_e_u*(1 – *x*)/2*N_e_x* ≈ Θ/(4*Ν_e_x*) for small x. This is proportional to the coalescence probability (~ 1/(2*N_e_x*)) in the same generation. Both coalescence and beneficial mutation typically occur very early in the selected phase when *x* is small. However, since their *relative* frequency is independent of *x*, their order along the genealogy is not affected. Consequently, mutation events will often happen on internal branches, leading to intermediate frequency haplotypes (Fig. 1C). In contrast, the probability of “picking” a recombinant ancestor, 2*N_e_x*(1 – *x*)*r*/2*N_e_x* = (1 – *x*)*r*, depends much weaker on *x*. During a sweep, recombination therefore tends to happen before mutation and coalescence (going back in time). It thus typically affects single external branches, leading to low-frequency polymorphism.

### Box: Inference of hard and soft sweeps

Can we identify patterns of soft sweeps? Clearly, only recent adaptive events leave a detectable footprint at all, hard or soft. Signals in the site frequency spectrum (like the excess of rare alleles that is picked up by Tajima’s D,Tajima, 1989) typically fade on time scales of ~ 0.1*N_e_* generations, while signals based on LD or haplotype statistics only last for ~ 0.01*N_e_* generations (Przeworski, 2002; Pennings and Hermisson, 2006b). For a clear footprint, selection must be strong (*4N_e_s_b_* ≫ 100). Even then, soft sweeps can be hard to distinguish from neutrality if they are “super soft”, i.e. if there are very many independent origins of the beneficial allele, or if its starting frequency in the SGV is high (≳ 20%, Peter et al., 2012; Berg and Coop, 2015).

For robust inference of selection against neutrality, we need a test statistic with consistently high power for hard and soft sweeps. As expected from the patterns described above, and as demonstrated (Pennings and Hermisson, 2006b; Ferrer-Admetlla et al., 2014), tests based on the site-frequency spectrum (looking for low- or high-frequency derived alleles) have low power to detect soft sweeps, while haplotype tests can detect both types of sweeps (Garud et al., 2015). In contrast to single-origin soft sweeps (which always leave a weaker footprint), the power to detect multiple-origin soft sweeps can be higher than the power to detect completed hard sweeps due to the conspicuous haplotype structure directly at the selected site (Pennings and Hermisson, 2006b).

Given that selection has been inferred, can we distinguish soft from hard sweeps? For soft sweeps with a single origin, this is difficult (Berg and Coop, 2015). Tests based on a combination of summary statistics have been developed by Peter et al. (2012) and by Schrider and Kern (2016a). Both tests have reasonable power to distinguish soft sweeps for strong selection and a high starting frequency (5 – 20%) of the selected allele. Clear empirical examples usually also rely on other evidence, complementing the footprint (Barrett and Schluter, 2008): e.g., a source population is known with the selected allele in the SGV (e.g. marine and freshwater sticklebacks,Colosimo et al., 2005), or a known and very recent selection pressure does not leave enough time for the allele to increase from a single copy to the frequency observed today (e.g. adaptation to HIV in humans, Novembre and Han, 2012).

Chances for a distinction of sweep types are better for soft sweeps with multiple origins. A tailor-made summary statistic, *H* 12, based on the two largest haplotype classes has been developed by Garud et al. (2015). It has a high power to distinguish hard and soft sweeps especially for recent or partial sweeps. *H* 12 has also been included into the deep learning algorithm by Sheehan and Song (2016) that is able to distinguish hard and soft sweeps.

## 3 Probabilities: When to expect soft sweeps

If adaptation is strictly mutation limited, all selective sweeps are necessarily hard. Without mutation limitation, soft sweeps can originate either from SGV or from recurrent new mutation (Fig. 2). Below, we discuss the probability for soft sweeps in both scenarios.

**Figure 2:**
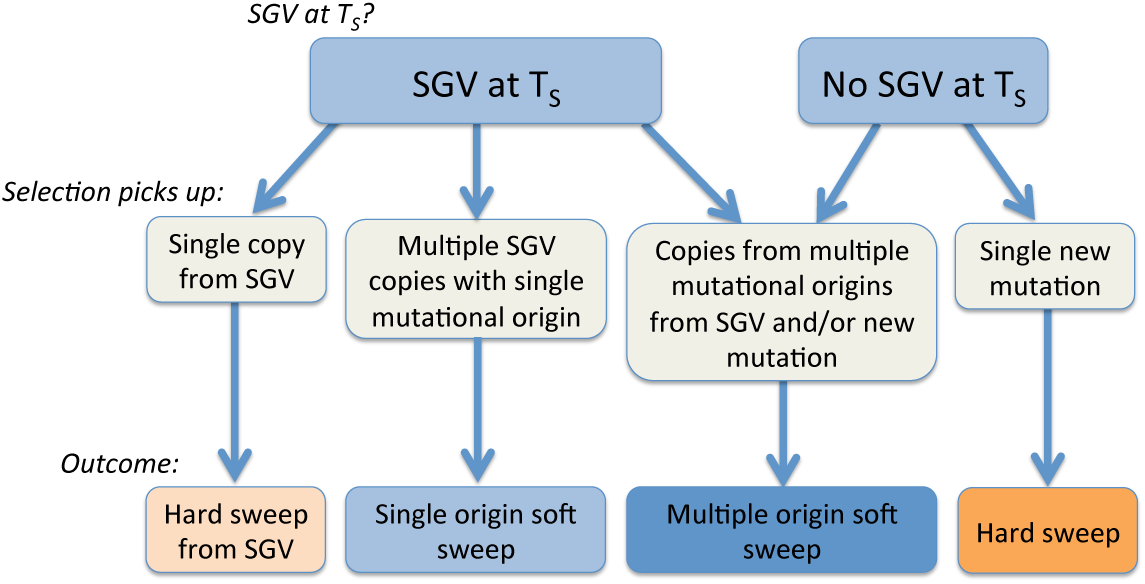
Possible sweeps types depending on whether SGV is available at the onset of selection (*T_S_*), and depending on how many copies or mutational origins of the beneficial allele are picked up by selection.

### Sweeps from standing genetic variation

When is adaptation from SGV likely? When does it produce hard or soft sweeps? In an early approach to address these questions, Orr and Betancourt (2001) used the following argument. Let *x*_0_ be the frequency of the A allele in the population at time *T_S_* when *A* turns beneficial. Assuming independence, each of the 2*N_e_x*_0_ mutant copies has the chance 2*s_b_* to escape stochastic loss. We can then approximate the distribution of the number *X* of *successful A* copies by a Poisson distribution with parameter *2s_b_*⋅2*N_e_x*_0_. The probability for a *standing sweep* (at least one successful mutant) is

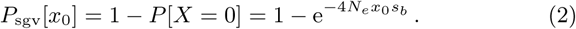

The conditional probability of a soft sweep (at least two successful copies), given that there is a standing sweep at all, follows as

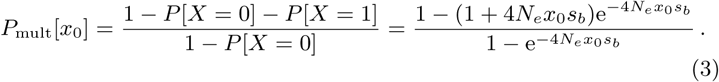

Clearly, *x*_0_ is a crucial parameter in these equations, but which value does it take? Orr and Betancourt (2001) and Jensen (2014) assume that the population is in mutation-selection balance prior to *T_S_* and use the deterministic approximation *x*_0_ = *u*/*s_d_* for the frequency of *A*. With Θ = 4*N_e_u*, we obtain 4*N_e_x*_0_*s_b_* = Θ*s_b_*/*s_d_*, which implies that hard sweeps (from SGV) dominate over soft sweeps (from SGV) as long as Θ*s_b_*/*s_d_* ≤ 1.14 (Orr and Betancourt, 2001). This implies, in particular, that hard sweeps from SGV always dominate for small Θ. However, as shown in Fig. 3A, this prediction is not correct.

**Figure 3:**
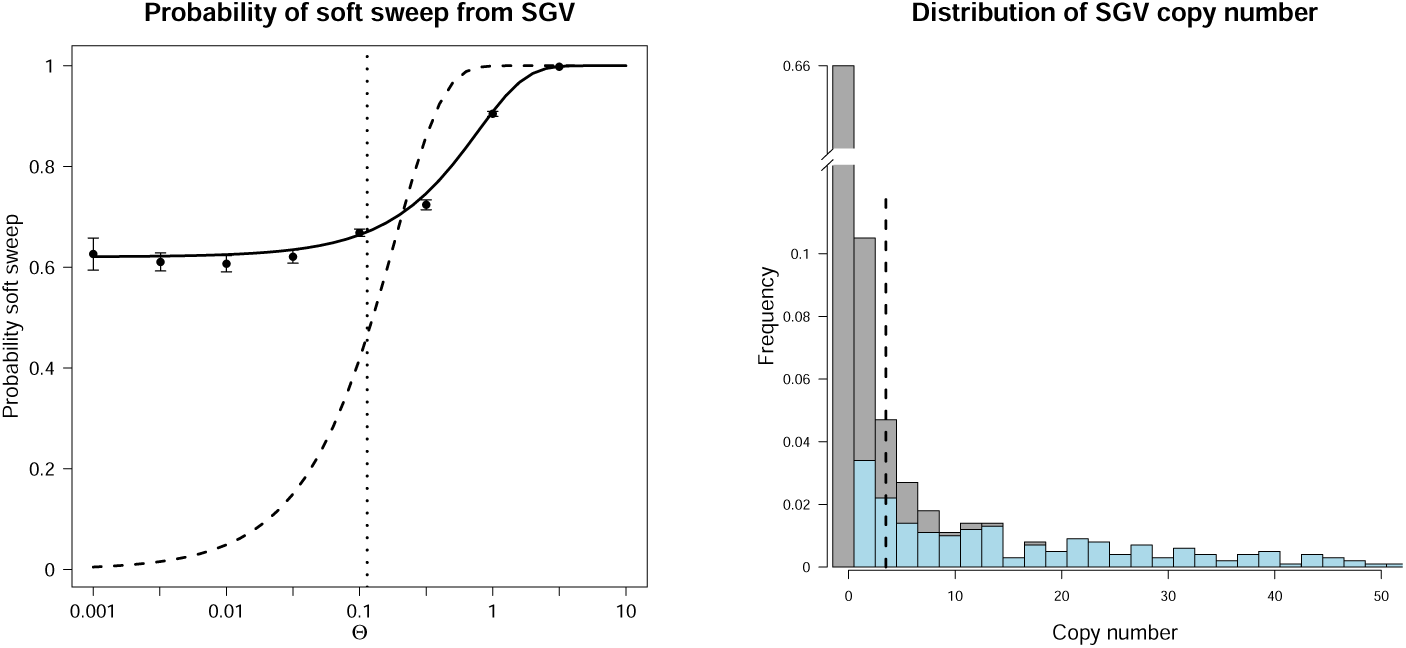
The probability that a sweep from SGV is soft. Panel A compares the deterministic (Eq. 6, dashed) and the stochastic approximation (Eq. 7, solid) to simulation results (dots with standard deviation). The deterministic approximation predicts that hard sweeps dominate for Θ < 0.114 (left of the vertical line). Panel B shows a simulated distribution for the number of copies of the beneficial allele in the SGV (grey) and the fraction that leads to a successful sweep (blue). The vertical dashed line marks the value *N_e_u/s_d_* **=** Θ*/*4*s_d_* assumed in the deterministic approximation. Parameters: *s_b_* = 0.1, *s_d_* = 0.01, Θ = 0.1 (in panel B).

The problem of the argument is that *x*_0_, the frequency of mutant allele A at time *T_S_*, is not a fixed value. Really, it is a stochastic variable and follows a distribution. As long as Θ < 1, this distribution is L-shaped with a maximum at 0. This means that in many cases, the *A* allele will not be present in the population, but in some cases, it will be present at a frequency that is much higher than *u*/*s_d_* (see figure 3B). Obviously, if the adaptive allele is not present in a population, a standing sweep cannot occur. On the other hand, if many copies of the allele are present, it is likely that a standing sweep will happen and that such sweep will be soft. The result is that, when the L-shaped distribution of *x*_0_ is taken into account, we get fewer sweeps in total, but those that do occur are more likely to be soft. We can compare how the two approaches predict the probability of a standing sweep (soft or hard) from mutation-selection balance. Using the deterministic assumption 4*N_e_x*_0_*s_b_* = Θ*s_b_*/*s_d_*, we find

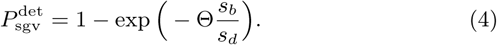

When taking into account the distribution of *x*_0_, we obtain (Hermisson and Pennings, 2005, Eq. 8)

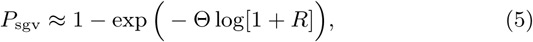

where *R* is the relative selective advantage defined in Eq. (1). Since log[1 + *R*] ≤ *R* < *s_b_*/*s_d_*, we have 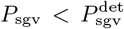. Next, we compare the two approaches as they predict the probability of a soft sweep, given that a standing sweep from mutation-selection balance has happened. The deterministic approximation reads

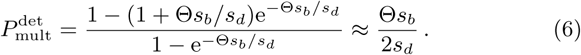

Accounting for the distribution of *x*_0_, we find (Hermisson and Pennings, 2005, Eq. 18)

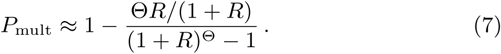

We can condition on the case of a single mutational origin of the allele by taking the limit Θ → 0, where (7) simplifies to

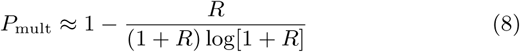

For R ≳ 4, we obtain more (single origin) soft sweeps than hard sweeps in the population, even for low Θ values, for which the deterministic approximation suggests that soft sweeps are exceedingly rare (Fig. 3A). For neutral standing variation (*s_d_* = 0), this requires only weak positive selection (*N_e_s_b_* > 1). For deleterious standing variation (*s_d_* ≫ 1/*N_e_*), we need *s_b_*/*s_d_* > 4. For biologically relevant parameters, this condition can deviate from the deterministic prediction by several orders of magnitude. For example, using the deterministic approximation, Jensen (2014) argues that soft sweeps are only likely if *s_b_*/*s_d_* > 100 when Θ = 10^−2^ (*“Drosophila”*) and if *s_b_*/*s_d_* > 10000 when Θ = 10^−4^ (“humans”) (Fig. 1 in Jensen, 2014). Our Fig. 3A shows that even for a low ratio *s_b_*/*s_d_* = 10, soft sweeps dominate for all Θ values.

### Sweeps from independent mutational origins

When will the genealogy of a beneficial allele contain multiple independent origins? A rough argument (Hermisson and Pennings, 2005; Messer and Petrov, 2013) shows that the probability of a multiple-origins soft sweep mainly depends on the mutation parameter Θ: A new beneficial allele establishes in the population within ~ log[4*N_e_s_b_*]/*s_b_* generations. During this time, further copies of the allele arise at rate ~ 2*N_e_u* and establish with probability 2*s_b_*. Establishment of a second beneficial mutation during the sweep thus becomes likely if 2*N_e_u*2*s_b_* log[4*N_e_s_b_*]/*s_b_* = Θlog[4*Ν*_e_*s_b_*] ⪆ 1. The basic message of this argument is correct. Detailed derivations based on coalescent theory (Pennings and Hermisson, 2006a) show an even weaker dependence on *s_b_*. For strong selection, 2*N_e_s_b_* ≫ 1, the probability for more than a single origin of the beneficial allele in a sample of size *n* derives to

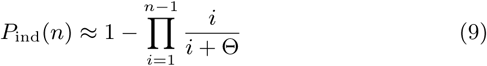

which simplifies to *P*_ind_(*n*) ≈ (0.577 + log[*n* – 1]) Θ for small Θ ≪ 1 and to *P*_ind_(*n*) ≈ 1 – (1/*n*) for Θ = 1.

Since Eq. (9) for *P*_ind_ depends only on Θ, but not on any selection parameter, it also applies if the selection pressure changes during the course of adaptation. In particular, the same result holds for adaptation from SGV (where selection changes from negative or neutral to positive) and for adaptation from recurrent new mutation after time *T_S_* only. Indeed, if Θ is large enough that adaptation acts on multiple origins of the same allele, the distinction of “standing variation” versus “new mutation” gets blurred – adaptive material is immediately available in both cases (Messer and Petrov, 2013). While the number of origins of a beneficial allele and their distribution in the sample are almost entirely independent of selection, other aspects of the sweep signature depend on selection strength. In particular, the signature is wide if and only if *s_b_* and *s_d_* are strong.

*P*_ind_(*n*) also depends only weakly on sample size *n*. This is because beneficial alleles from different origins are typically at intermediate frequencies in the population. Indeed, a multiple-origin soft sweep in the entire (panmictic) population will usually be visible already in a sample of moderate size. For *n* = 100, multiple-origin soft sweeps start to appear for Θ > 0.01 (> 5% soft), become frequent for Θ > 0.1 (≈ 40% soft) and dominate for Θ ≥ 1 (≥ 99% soft).

The probabilities of adaptation from SGV (5) and from multiple mutational origins (9) are both governed by the population mutation parameter Θ = 4*N_e_u*. Before we can assess Θ in natural populations, however, the factors that enter this composite parameter require some elaboration.

### Understanding *N_e_* in Θ: the effective population size

We have, so far, not made any distinction between the census size of a population and its so-called effective size. For a better understanding, we need to do this now. The expected number of new copies of a mutant allele A that enters a diploid population each generation is twice its census size times the mutation rate, 2Nu. For species as abundant as microbes, fruitflies, or humans, almost any adaptive mutation that is possible will appear many times within a short time interval. However, what matters for adaptation are only those mutants that escape genetic drift. The establishment probability of a mutant with selection coefficient *s_b_* is roughly 2*s_b_N_e_*/*N*. Here, the effective population size *N_e_* is an (inverse) measure of the strength of genetic drift: strong drift (small *N_e_*) leads to a higher chance of loss due to random fluctuations. We then see that the rate of *successful* mutants, 2*Nu*⋅*2s_b_N_e_*/*N* = 4*N_e_us_b_*, depends only on *N_e_*, while the census size drops out.

How should we determine *N_e_*? Many factors influence the strength of genetic drift in natural populations (Charlesworth, 2009). Some factors, such as variance in offspring number or unequal sex-ratios, act on the population in every single generation; others recur every few generations, like seasonal fluctuations of the census size. If all relevant factors operate on shorter time scales than typical coalescence times, they can be subsumed in a single well-defined *coalescent effective population size N_e_* (Sjödin et al., 2005). Since, by definition, the coalescent effective size is constant over time, it can readily be estimated from neutral polymorphism or diversity data using measures like Watterson’s Θ*w*.

Unfortunately, several factors that drive drift do not fit this scheme. These include fluctuations in population size over time scales comparable with coalescence times and non-recurrent/sporadic events, such as ongoing population growth, single population bottlenecks, or episodes of linked selection. If any such event occurs during the coalescent history of an allele, it exerts a drift effect on its frequency distribution, but there is “no meaningful effective population size” (Sjödin et al., 2005) to fully describe this effect. Therefore, the question arises which value for *N_e_* should be used in the parameter Θ = 4*N_e_u* that enters our formulas.

Even if it does not take a unique value, *N_e_* is always a measure of drift. In each particular case, we therefore should ask which aspect of drift is relevant. For example, equilibrium levels of SGV are affected by drift throughout the coalescent history of potential “standing” alleles. If adaptation occurs from neutral variation, this history is long and a long-term measure of genetic drift, such as neutral polymorphism Θ*w* or pairwise nucleotide diversity *π* can serve as a valid proxy for Θ in formulas like Eq. (5).

For the probability of adaptation from multiple independent origins (9), the decisive time period where drift is relevant is only the establishment phase of the beneficial allele (Pennings and Hermisson, 2006a). This is a time window of length ~ 1/*s_b_*, where a successful allele quickly spreads. For very strong selection, this period can be as short as 10 – 100 generations. We thus need an estimate for a “fixation effective size” (Otto and Whitlock, 1997) or “short-term *N_e_*” (Karasov et al., 2010; Barton, 2010), which can differ from the “long-term *N_e_*”: While factors like the offspring variance enter also into a short-term *N_e_*, any event that occurs outside of the crucial time window has no effect. Unfortunately, there is no easy direct measure of the short-term *N_e_* from polymorphism data. In particular, if demographic factors are at play, using neutral diversity can lead to great underestimates of *N_e_*. To account for such major demographic trends, one can use inference methods like PSMC (Li and Durbin, 2011) or deep sequencing (e.g. Chen et al., 2015) to estimate a time-dependent effective size *N_e_*(*T_S_*) at the (putative) time of the adaptation. However, due to limited resolution, these methods will never capture all confounding demographic factors. As pointed out by Karasov et al. (2010), the same holds true for effects of linked selection (recurrent sweeps). Since unresolved factors usually (though not always) increase drift, estimates of *N_e_*(*Ts*) generally still underestimate the “true” short-term *N_e_*.

Values of short-term *N_e_* vary not only between populations, but also across adaptive loci along the genome. They depend on the exact timing of the selection windows relative to demographic events or episodes of linked selection. Genome-wide studies, e.g. in *D. melanogaster,* show high heterogeneity in diversity levels due to linked selection (Elyashiv et al., 2016). Finally, since stronger selection leads to shorter windows, the relevant short-term *N_e_* may also depend on the selection strength (Otto and Whitlock, 1997). As Wilson et al. (2014) point out, this may lead to larger *N_e_* (thus larger Θ and more soft sweeps) for strong adaptations with shorter establishment times.

### Understanding *u* in Θ: the allelic mutation rate

For any adaptive mutant allele *A*, e.g. a resistant phenotype, we can ask how this allele can be produced from a wildtype by mutation of the underlying molecular genotype. Sometimes, *A* is highly specific and can only be generated by a unique (point-)mutation. However, often multiple mutations produce the same phenotype (e.g. Barroso-Batista et al., 2014, for *E. coli* adaptation in the mouse gut). This is a generic property of the genotype-to-phenotype map, which often maps whole classes of equivalent genotypes to the phenotype that is seen by selection. Redundancies already exist on the level of the genetic code, but also on any other level, both in the coding sequences or regulatory regions of single genes, and across genes and pathways. In this case, *A* has an extended *mutational target* (Pritchard et al., 2010), which is reflected by an increased allelic mutation rate *u.* For strict redundancy, we require that all mutations in the target of *A* have the same fitness effect. We further require that multiple redundant mutations do not increase fitness any further.

We need to distinguish mutational targets on two different levels. On the level of a single locus, *u_l_* and Θ_*l*_ = 4*N_e_u_l_* measure the total rate of redundant mutations to produce allele *A* in a single recombinational unit (technically, we need that recombination during the selective phase is unlikely). Mutations contributing to Θ*l* interfere both due to complete linkage (as in clonal interference) and due to epistasis. Without epistasis, the dynamics of adaptation under recurrent mutation is driven by *mutation stacking* and is dominated by the haplotype most loaded with beneficial mutations (Desai and Fisher, 2007). Negative epistasis among redundant mutations erases the fitness advantage of stacking and fundamentally changes the dynamics of adaptation, leading to soft sweeps instead of clonal interference. (We note that our definition of a mutational target deviates from the one by Messer and Petrov (2013), who also include adaptations to unrelated selection pressures.)

On the genome level, *u_g_* and Θ_*g*_ = 4*N_e_u_g_* cover all redundant mutations that produce an *A* phenotype across all loci. Thus, Θ_*g*_ = *m*Θ_*l*_ for *m* equivalent loci. Mutations for *A* on different genes may be unlinked, but still interfere due to negative epistasis. Without epistasis, the dynamics across different loci decouples, reproducing hard or soft single-locus sweeps. With epistasis, patterns and probabilities are no longer independent and new phenomena can arise. Currently, the corresponding patterns of polygenic adaptation (Pritchard et al., 2010) that can be distributed across many loci remain largely unexplored (see Berg and Coop, 2014, for the case without epistasis). However, simulations show that often a single locus emerges that contributes most to the effect. In our simulations below, we study how a multi-locus target Θ_*g*_ > Θ_*l*_ affects the single-locus probabilities of hard *vs.* soft sweeps at this focal locus.

Θ_*l*_ and Θ_*g*_ affect the probabilities of hard and soft sweeps in different ways. The probability of a multiple-origin soft sweep, Eq. (9), depends only on the mutation rate Θ_*l*_ of the corresponding locus. In contrast, the probability for a single-origin soft sweep *vs.* a hard sweep depends primarily on the genome-wide rate Θ_*g*_. Indeed, the probability of adaptation from SGV depends on whether some of this variation, genome-wide, is picked up by selection. Therefore, Θ_*g*_ should be used in Eq. (5). If adaptation happens from SGV, the *conditional* probability for adaptation from multiple SGV copies at a locus, Eq. (7), depends on Θ_*l*_, but becomes independent of Θ_*l*_ in the limit of a single mutational origin (Eq. 8). Although hard and soft sweeps are single-locus footprints (defined via a single-locus genealogy), their probabilities thus depend on both Θ_*l*_ and Θ_*g*_. Note that a short-term *N_e_* is relevant for recurrent mutation and thus Θ_*l*_, while a long-term *N_e_* is relevant for total levels of variation and thus Θ_*g*_.

Depending on the nature of the adaptation, allelic mutation rates *u_i_* and *u_g_* can vary widely. On the low end of the scale are phenotypes that are only generated by a single base substitution. In this case, *u_g_* = *u_l_* can be as low as *μ*/3, where *μ* is the point mutation rate (assuming that all three base substitutions occur at equal rates and ignoring variance of *μ* along the genome). On the locus level, mutational targets can comprise 10s and maybe 100s of point mutations (and also include insertions/deletions). High values for *u_l_* should be expected especially for adaptive loss-of-function mutants and for some (cis-) regulatory mutations, as in the case of the Lactase gene (see below). For alleles with a polygenic target, *u_g_* could in turn be an order of magnitude larger than the locus rate *u_l_*.

It is worth noting that neither *u_l_* nor *u_g_* are closely related to the so-called distribution of fitness effects (DFE), which, among other things, informs us about the proportion of beneficial mutations among all mutations that hit some target (Jensen, 2014). The DFE divides the total mutation rate into classes of deleterious, neutral, and beneficial and generally finds that beneficial mutations are only a small fraction. In contrast, *u_l_* and *u_g_* ask for the total mutation rate to specific beneficial allele *A* of which we know that it exists. As such, it is not affected by the presence of further, neutral or deleterious mutations on the same target.

### Simulation results

Figure 4 shows percentages of soft and hard sweeps for beneficial alleles with single-locus or multiple-locus mutation targets and with strong or weak fitness trade-off in the ancestral environment. For soft sweeps, we distinguish single- and multiple-origin types. For hard sweeps, we specifically record cases that derive from a single ancestor in the SGV. Assume first that adaptation is locus-specific and can only occur at a single gene, Θ_*g*_ = Θ_*l*_ (Fig. 4, top row). This may often be the case for resistance mutations.

- For mutations with a strong trade-off, *s_d_ ≥ s_b_*, hard sweeps dominate for Θ_*l*_ < 0.1, while multiple-origin soft are most prevalent for Θ_*l*_ > 0.1. Adaptations from SGV are only likely for Θ_*l*_ ≫ 0.1 where they produce multiple-origin sweeps. Single-origin soft sweeps or hard sweeps from SGV are very rare already for *s_d_* = *s_b_*, R ≈ 1 (Fig. 4A). For adaptations with even stronger trade-off (e.g. *s_d_* = 5*s_b_*, as suggested by Orr and Betancourt, 2001; Jensen, 2014), they are virtually absent.
- For mutations with a weak trade-off, *s_d_* ≪ *s_b_*, the probability for adaptation from SGV is higher (cf Fig. 4B with R ≈ 100). We now find a few single-origin soft sweeps for intermediate Θ values, while hard sweeps from SGV remain rare.

**Figure 4:**
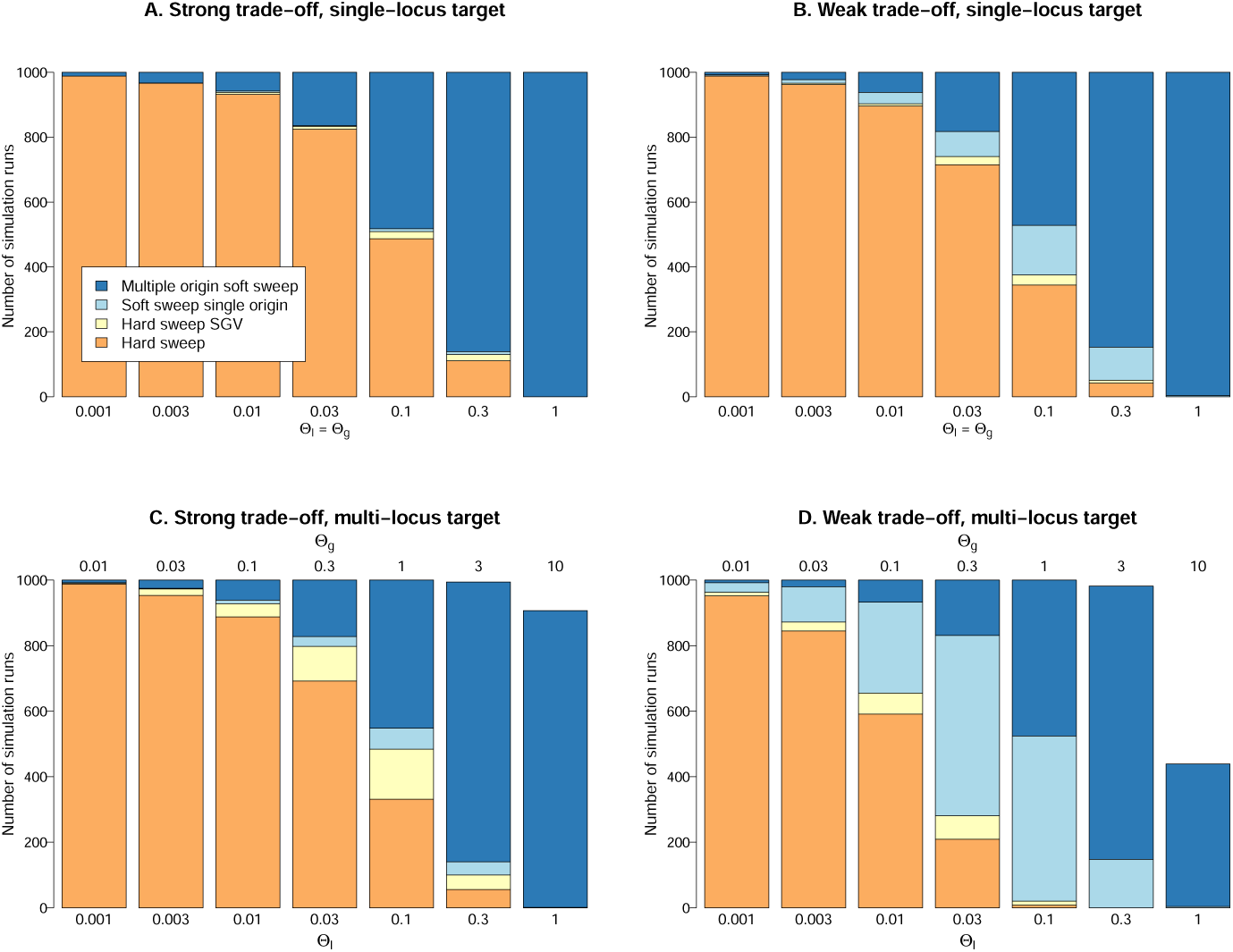
The probability of hard and soft sweeps for beneficial alleles with single-locus mutation target (top row) and for alleles with a target of 10 identical loci (bottom row). 2*N_e_* = 10000 and *s_b_* =0.1. Panels A and C assume a strong fitness trade-off *s_d_* = *s_b_* = 0.1. Panels on the right side assume a weak trade-off *s_d_* = 0.001. See the Methods section for further details on the simulations.

The single locus results also describe adaptation of a polygenic trait as long as mutations at different loci do not interact. This is different if the beneficial allele has a mutational target across several loci, Θ_*g*_ > Θ_*l*_, where mutations interact by negative epistasis. Fig. 4, (bottom row) shows how this affects probabilities of soft and hard sweeps, assuming 10 identical loci, Θ_*g*_ = 10Θ_*l*_. Although adaptation can occur, in principle, by small frequency shifts at many loci, we usually observe a *major sweep locus* that experiences a frequency shift of > 50%. For small Θ_*g*_, adaptation is often even entirely due to a single locus. The figure shows the sweep type at a major sweep locus and leaves all areas white where no such locus exists.

- The effect of a multi-locus target is an increase in the probability of adaptation from SGV (Fig. 4C/D). Hard sweeps from new mutations are pushed back to lower values of Θ_*l*_. In a parameter range with intermediate Θ_*l*_ between 0.01 and 0.1, hard sweeps or single-origin soft sweeps from SGV can occur. For a strong trade-off (Fig. 4C), these sweeps still produce essentially hard sweep patterns. We thus obtain the same distribution of patterns as in the single-locus case (compare with Fig. 4A).
- The combination of a multi-locus target and a weak trade-off creates the largest parameter space for adaptation from SGV (Fig. 4D). For s_b_ ≫ *s_d_*, almost all sweeps from SGV are soft (single- or multiple origin). The expected footprint of all these sweeps is clearly distinct from a hard sweep.
- Finally, for high mutation rates Θ_9_ ≳ 3, we obtain cases of polygenic adaptation with small frequency shifts at many loci or super-soft sweeps with very large starting allele frequency in the SGV. In these cases, no major sweep locus exists (white areas in the figure).

Across all cases, the probability for a multiple-origin soft sweep depends only on the locus mutation parameter Θ_*l*_ but not on Θ_*g*_. It also does not dependent on selection strengths *s_b_* and *s_d_*, on epistasis among loci, or on the presence or absence of SGV. In contrast, the probabilities for hard sweeps and single-origin soft sweeps depend on on both, Θ_*l*_ and Θ_*g*_, and on selection strength. For adaptations with a single-locus target, Θ_*g*_ = Θ_*l*_, or strong trade-off, *s_d_* ≥ *s_b_*, both of these sweep types are rare (Fig. 4A-C). This only changes for beneficial alleles with a multi-locus mutation target, Θ_*g*_ ≫ Θ_*l*_ and weak trade-off, *s_d_* ≪ *s_b_*, where single-origin soft sweeps dominate for intermediate Θ_*l*_ values (Fig. 4D).

Our analysis of the multi-locus case is necessarily limited. While we assume full redundancy, mutations at different loci are often only partially redundant. They can have variable fitness effects and negative epistasis (which is a natural consequence of stabilizing selection) can be weaker. It remains to be explored how these factors influence the probabilities of sweep types at single loci, as well as the tendency for polygenic footprints with small allele frequency changes at many loci. Another factor that is ignored here, but which increases the percentage of sweeps from SGV among all observed sweeps comes to play if we condition on a short time period between the onset of selection and the time of observation (Hermisson and Pennings, 2005; Pritchard et al., 2010; Berg and Coop, 2015). Further complications, e.g. due to population structure, are briefly discussed below. Any comprehensive analysis of the probabilities of sweep types in genome scans needs to take these factors into account.

## 4 Complications

### Population structure

Most natural populations are spatially extended and show at least some degree of geographic structure. While our genealogy-based definition of sweep types applies in the same way, the probabilities to observe hard or soft sweeps can depend on the strength of structure and also on the sampling strategy. Consider a spatially structured population that experiences a common novel selection pressure. If genetic exchange between different parts is weak, adaptation across the whole range may require independent origins of the adaptive allele in different geographic regions, a phenomenon also called *parallel adaptation* (Ralph and Coop, 2010). Thus, spatial structure can increase the probability of multiple-origin soft sweeps in global population samples, but not necessarily in local samples from a single region.

#### When does population structure favor soft sweeps

If migration is long-ranged (like in an island model), only very strong structure (weak migration) will increase the probability of a soft sweep in global samples (Messer and Petrov, 2013). Consider two islands of size *N_e_*, each with Θ = 4*N_e_u* ≤ 0.01 (assuming Θ = Θ_*l*_ = Θ_*g*_), such that soft sweeps within islands are unlikely. In this regime, an adaptive mutation will fix on one island before an independent mutation can establish on the other island. If migration is larger than mutation, 4*N_e_m* > Θ, it is more likely that the second island adapts from a beneficial migrant than from independent mutation, which results in a hard sweep. Soft sweeps are only likely if 4*N_e_m* < Θ (less than a single migrant per 100 generations) (Messer and Petrov, 2013). Note however, that effective isolation is only required during a short time window. This could increase the probability of spatial soft sweeps during periods of fragmentation, e.g. in glaciation refugia.

If migration is short-ranged and causes isolation by distance, geographic structure has a stronger effect (Ralph and Coop, 2010, 2015a). Consider a population of size *N_e_* in continuous space with average dispersal distance *σ*. In such models, adaptations spread from their point of origin in so-called Fisher waves that proceed with constant speed 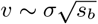. In populations that extend over *d* dispersal distances, the time-lag due to the finite wave speed enables parallel adaptation with multiple waves if 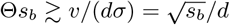 (Ralph and Coop, 2010; Messer and Petrov, 2013). If parallel waves are driven by different origins of the same beneficial allele at the same locus, they constitute a spatial soft sweep. With large diameter *d*, spatial soft sweeps can occur for lower Θ than multiple-origin soft sweeps in the panmictic case (9), especially for strong selection (large *s_b_*) or for alleles with weak trade-off (small *s_d_*) that adapt from SGV (cf Ralph and Coop, 2015a).

#### Patterns of spatial soft sweeps

Whenever spatial structure and isolation by distance cause soft sweeps, despite low Θ, we expect to find hard sweeps in local samples, but soft sweeps in global samples. However, as Ralph and Coop (2010) observe, ongoing migration can blur this signal and patterns can look increasingly soft in local samples, too.

Vice-versa, there is also a parameter range where population structure is not causal for multiple origins, but still leads to geographic sweep patterns. Consider a population with Θ ~ 1, such that multiple-origin soft sweeps are likely. Now, assume that this population is divided into *k* islands. We still have multiple origins globally, but if Θ/*k* ≪ 1 multiple origins on a single island are unlikely. Assume that islands are connected by migration of strength 1 ≪ 4*N_e_m* ≪ 2*N_e_s_b_.* In that case, gene flow will erase any trace of structure at neutral sites. However, migration during the selection window of ~ 1/*s_b_* generations is limited. Samples taken from an island where a beneficial mutation has occurred will thus be dominated by descendants from this mutation and may show patterns of local hard sweeps.

In natural populations, further factors can influence probabilities and patterns of soft sweeps. For example, heterogeneous spatial selection promotes parallel adaptation if gene flow between “adaptive pockets” is hampered by negative selection in interjacent regions (Ralph and Coop, 2015b). In contrast, gene surfing during range expansions may hamper soft sweeps, because successful copies of the beneficial allele need to originate in a narrow region near the expansion front (Gralka et al., 2016). Currently, patterns of spatial soft sweeps have only partially been characterized and remain an active field for study.

### Soft sweeps turning hard?

Sweeps that appear soft for a recent adaptation may turn hard if we take a sample at some later time. This occurs if descendants from only a single copy of the beneficial allele dominate at that later time, either because of genetic drift or due to ongoing selection.

Under neutrality, the average time to fixation or loss of an allele at frequency *x* is ≈ 4*N_e_*(*x* log *x*^−1^ + (1 – *x*) log(1 – *x*)^−1^) generations (Ewens, 2004). Soft sweeps typically lead to intermediate haplotype frequencies, but even for a major haplotype frequency of 99% the expected fixation time is > 0.1*N_e_* generations. I.e., over time scales where sweep patterns are generally visible, a soft sweep will usually not “harden”. Demographic events like bottlenecks increase drift and lead to accelerated hardening. Still, a single bottleneck needs to be very strong to erase a soft sweep pattern. By genetic drift alone, soft sweep signals will rather fade than turn hard.

Fitness differences among soft sweep haplotypes can occur either because different mutations in the target of the beneficial allele are not fully redundant or because of selection on linked variation. As long as these differences are small relative to the primary effect of the allele, patterns of recent soft sweeps appear to be remarkably stable. Pennings and Hermisson (2006a) show that the patterns and probabilities of multiple-origin soft sweeps do not change much for fitness differences up to 50% of the primary effect if the sample is taken at fixation of the primary allele. However, this no longer holds if fitness differences are of the same order as the primary effect or if sampling occurs at a later time. For fitness differences of 2*N_e_*∆*s_b_* ≥ 100, hardening occurs within the time window of ~ 0.1*N_e_* generations where selection footprints are visible. An example of such hardening in *Plasmodium* is described in the section on Microbes. A related issue arises if a locus is hit by recurrent sweeps. Indeed, both a stepwise approach of a new optimum and compensatory mutation at the same gene are well-documented adaptive responses to a new selection pressure. If adaptation is not mutation limited, these steps can occur in quick succession. Due to such “stacking” of soft sweeps (repeated Ewens sampling), we readily obtain a hardening of the pattern, unless the locus mutation rate is as high as Θ_*l*_ ≥ 1, where many haplotypes survive each sweep.

Importantly, a hardened soft sweep is not the same as a classical hard sweep, neither its biological interpretation (see the *Plasmodium* example below), nor its pattern. Both stages of the process are visible in time-series data and potentially also in the final pattern, e.g., if a strong (soft) resistance adaptation is followed by a weaker (hardening) compensatory mutation at the same gene. Theory and statistical methods to deal with such complex cases are currently lacking.

### Hard sweeps looking soft?

In a recent paper, Schrider et al. (2015) suggested that patterns that look like soft sweeps could result from flanking regions of hard sweeps (soft shoulder effect). It is easy to see what could drive a spurious signal: for many summary statistics of polymorphism data, like *π* or the number of haplotypes, both, soft sweeps and flanking regions of hard sweeps result in weaker footprints than the core regions of hard sweeps. Selection scanners that base their prediction on a single local window can confuse one for the other. However, Schrider et al. (2015) do not provide an example where a pattern has been misclassified because of the shoulder effect. It seems that problems can be avoided if scans pre-select loci with strong signals (Garud et al., 2015; Garud and Petrov, 2016) or if selection footprints are evaluated together with their local genomic context (Sheehan and Song, 2016; Schrider and Kern, 2016a).

Another concern that has been voiced is that allelic gene conversion during a hard sweep could mimic a soft sweep pattern (Schrider et al., 2015). Indeed, gene conversion at the selected site can change the pattern of a hard sweep (Pennings and Hermisson, 2006b; Jones and Wakeley, 2008): A conversion event with short conversion tract around the selected allele can place this allele onto another genetic background. Like for a multiple-origin soft sweep, the beneficial allele becomes associated with several ancestral haplotypes that stretch across the adaptive target. LD across the selected site will be positive, in contrast to a classical hard sweep, and in accordance with a soft sweep. However, important differences remain. Like single crossing-over, gene conversion is a recombination event. As explained in the Footprints section, recombination during a hard sweep typically leads to recombination haplotypes at a low frequency in the sample (see also Jones and Wakeley, 2008), in contrast to the intermediate-frequency haplotypes created by recurrent mutation at a soft sweep (Pennings and Hermisson, 2006b).

We can apply this argument to the polymorphism pattern in three immunity genes in *Drosophila simulans* reported by Schlenke and Begun (2005). All three genes show extreme values of positive LD, caused by invariant haplotypes that extend across the gene. In each case, two haplotypes are found at an intermediate frequency in a combined Californian sample. Whereas it appears unlikely that such a pattern, repeated across three genes, has been created by gene conversion, it is perfectly expected under the scenario of multiple-origin soft sweeps. Clearly, gene conversion can also be excluded if different redundant mutations at the same locus contribute to a soft sweep pattern (as for the lactase case discussed below).

A case where a hard sweep will look soft results if selection acts on a fully recessive allele. A recessive allele behaves essentially neutrally as long as its frequency is smaller than 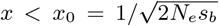. Its trajectory resembles the one of an allele that derives from neutral SGV with starting frequency *x*_0_ at time *T_S_*. Since the impact of selection on the sweep depends only on the shape of the trajectory, both footprints are indistinguishable. Indeed, equivalent models have been used to describe recessive hard sweeps (Ewing et al., 2011) and single-origin soft sweeps from SGV (Berg and Coop, 2015). As stressed by Berg and Coop (2015), both scenarios can only be distinguished if additional biological information about dominance is available. Note, finally, that the combined effect of a recessive hard sweep or single-origin soft sweep *and* gene conversion may indeed create a pattern that mimics a multiple-origin soft sweep.

## 5 Evidence

How do our theoretical expectations of sweep types compare with observed data? Evidence for both soft and hard sweeps exist for many species. In this section, we focus on three exemplary cases: *Drosophila,* humans, and microbial adaptation.

### Drosophila

Estimates of Watterson’s Θ*w* per base pair from the least constrained regions (short introns) in African *D. melanogaster* and *D. simulans* (Parsch et al., 2010; Andolfatto et al., 2010) result in lower bounds Θ_*l*_ ≥ Θ_*w*_/3 ≈ 0.008 (for *mel*), and Θ_*l*_ ⪆ 0.013 (for *sim*), respectively. These values are right at the boundary where multiple-origin soft sweeps start to appear in larger samples (4% resp. 6% in a sample of size 100, Eq. 9). Given that short-term N_e_ likely exceeds the values from polymorphism data and mutational targets may often consist of at least a few sites, Θ_*l*_ ~ 0.1 is a plausible estimate for the per-locus rate. We thus expect to see a mixture of hard sweeps and multiple-origin soft sweeps in data (Fig. 4). For adaptations with a weak trade-off, also single-locus soft sweeps can contribute, in particular if mutation targets are polygenic (Θ_*g*_ > Θ_*l*_).

Recent genome scans by Garud and Petrov (2016) and Sheehan and Song (2016) for a Zambian *D. melanogaster* population confirm to this expectation. Both find a mix of hard and soft sweep signals for regions with the strongest evidence for recent selection. Garud et al. (2015) and Garud and Petrov (2016) report much higher rates for soft sweeps for *melanogaster* from North Carolina. According to their test, all top 50 signals are consistent with soft sweeps rather than hard sweeps. Where this difference comes from is yet unclear. Potential causes include more recent selection in the American population and a larger short-term *N_e_*, but also confounding factors like inbreeding and admixture.

Similar to genome scans, also an inspection of selection footprints at genes with well-characterized function shows a mix of signals. However, clear patterns that are fully consistent with the simplest sweep models are rather the exception. Among the clearest examples are patterns consistent with multiple-origin soft sweeps in *D. simulans* immunity receptor genes (Schlenke and Begun, 2005, discussed above) and the loss of pigmentation in D. *santomea* due to inactivation of a cis-regulatory element at the *tan* locus (Jeong et al., 2008). The latter case is an example for an adaptive loss-of-function allele that can be produced by a large number of fully redundant mutations (three independent origins have been identified, two deletions and one double substitution). The adaptation underlies a species characteristic of D. *santomea* and is a rare case of a sweep (hard or soft) with clear phenotypic effect that is completed throughout the species.

A well-resolved example for a partial hard sweep is the *Bari-Jheh* insertion in *D. melanogaster,* a gain of function mutation for protection to oxidative stress (Gonzáles et al., 2008; Guio et al., 2014). Since the allele has a large fitness trade-off, it is unclear whether it is *de-novo* or from SGV. Both cases would result in a hard sweep pattern. An example for a partial sweep from SGV is the *CHKov1* gene in *D. melangaster* (Aminetzach et al., 2005). At the gene, an exonic insertion that provides insecticide resistance has recently swept to high frequency. Strong divergence of the sweep haplotype shows that the allele must be very old and long predates the selection pressure. However, the pattern is not easily explained by a simple selection history. Indeed, the allele most likely has a history of pre-adaptation to viral infection (Magwire et al., 2011) and the signal is shaped by multiple selection episodes. Note that single-origin sweeps (soft or hard) are likely for exonic insertion adaptations, which are not easily replicated and plausibly have a very low allelic mutation rate.

A stepwise selection history is also evident for evolution of resistance to oranophosphate insecticides at the *Ace* locus in *D. melanogaster* (Menozzi et al., 2004; Karasov et al., 2010; Messer and Petrov, 2013). In response to a very recent selection pressure, the same base substitution has occurred at least three times independently in different world regions (global soft sweep) and partially also within regions (local soft sweep). Resistance is reinforced by two further substitutions at the *Ace* locus. The haplotype pattern of the most resistant type shows signs of partial (but not complete) hardening.

### Humans

For Humans, estimates of a long-term *N_e_* ~ 10^4^ (Takahata, 1993) are so heavily influenced by sporadic demographic events, like the bottleneck connected to out-of-Africa migration, that this number is almost useless for population genetic theory. More refined methods based on deep sequencing estimate changes in *N_e_* from ~ 14000 pre-agriculture (10000 years ago) to ~ 500000 presently in Africa (Chen et al., 2015). With a mutation rate of 1 – 210^−8^, this leads to a “recent” Θ ≈ 10^−3^ for point mutations, consistent with estimates of Θ from singletons (Mathieson and McVean, 2014). Using similar assumptions about target sizes and shortterm N_e_ as for *Drosophila,* we arrive at ~ 0.01 as a rough estimate for Θ_*l*_ or Θ_*g*_. This is an order of magnitude lower than for *Drosophila* and in a range where hard sweeps from new mutations dominate (Fig. 4). However, there are reasons why adaptation from SGV may be more prevalent. First, population growth protects rare alleles from loss due to drift (Otto and Whitlock, 1997; Hermisson and Pennings, 2005). This enhances the probability of adaptation from SGV and is not captured by Eq. (5). Second, many selection pressures result from the dramatic changes in nutrition and population density since the advent of agriculture, or from pathogens that have spread in response to increased human density. Since these selection pressures are very young, almost all adaptive alleles that are now at high frequency must have emerged from SGV (Przeworski et al., 2005; Novembre and Han, 2012). Depending on their fitness trade-off or potential pre-adaptation in the ancestral environment, adaptations will display signals of either hard or soft partial sweeps.

Tests for candidate genes and genome scans that can distinguish hard and soft sweeps either find evidence for both types (Peter et al., 2012; Schrider and Kern, 2016a) or predominantly soft sweeps (Schrider and Kern, 2016b). Detailed studies of individual genes support the role of SGV in recent adaptations. A clear example is the CCR5 mutation conferring resistance against HIV (reviewed by Novembre and Han, 2012). Other loci once again reveal more complex adaptive histories. E.g., the recent spread of a derived haplotype at a Serine protease inhibitor gene in Yorubans has been identified as resurrection of a pseudogene using pre-adapted SGV (Seixas et al., 2012). Adaptation to high altitude in Tibetans involves a variant of the hypoxia pathway gene EPAS1 that has introgressed into modern humans from Denisovans (Huerta-Sanchez et al., 2014).

A consistent finding is that recent sweeps in humans are never species-wide (Mallick et al., 2016), but rather partial and regional. Regional patterns arise not only as a response to regional selection pressures (e.g., adaptation to Lassa fever in regions where the disease is endemic, Andersen et al., 2012), but also because of parallel adaptation to the same selection pressure across geographic regions (Coop et al., 2009; Ralph and Coop, 2010, 2015a; Novembre and Han, 2012). The prime example is light skin pigmentation, a trait that has evolved several times independently in Europeans and Asians. Parallel adaptation in this case is facilitated by a larger genomic mutation target, comprising several genes. The pattern at each single gene locus is still consistent with a hard sweep, either from new mutation after the out-of-Africa migration or from a deleterious standing variant (Coop et al., 2009; Jablonski and Chaplin, 2012).

Given the low estimated value of Θ_*l*_, maybe the most surprising finding is the evidence of multiple-origin soft sweeps – including two of the most prominent examples of recent adaptation in humans: Besides the lactase adaptation, which has a large mutation target (see separate box), this is the *Duffy* FY-0 mutation conveying resistance against Vivax malaria. For *Duffy,* the exact same point mutation has been found on three different haplotypes. Two haplotypes with linkage disequilibrium across the selected site are found at intermediate frequency in sub-Saharan Africa (Hamblin and Di Rienzo, 2000). One further independent origin has been described in Papua New Guinea (Zimmerman et al., 1999). There is evidence for the influence of geographic structure for both, lactase and *Duffy* (global soft sweep), but both are also locally soft in Africa.

### Box: The human lactase gene example

The lactase enzyme enables humans (and other mammals) to digest the milk sugar lactose and to consume milk without adverse side-effects. All humans produce lactase as infants, but in the ancestral wildtype the gene is down-regulated during childhood. However, around 61% of Europeans, 34% of Africans, and 28% of East Asians today are lactose tolerant also as adults (Ingram et al., 2009a). The trait has been attributed to derived allelic variants in an enhancer region of LCT (the lactase gene). To date, 5 different functionally confirmed mutations have been identified, all in a narrow region of around 100bp, 14000bp upstream of LCT (Tishkoff et al., 2007; Ingram et al., 2009a; Jones et al., 2013). They display clear geographic structure: lactose tolerance in Europe and central Asia is almost exclusively due to a single SNP, while a different causal SNP dominates among milk consumers in Tanzania and Kenya. Polymorphism data from LCT in these populations show haplotype patterns compatible with a strong partial sweep (Bersaglieri et al., 2004; Tishkoff et al., 2007) that is globally soft, but hard in each single region. However, more recent studies in Ethiopian and Sudanese lactose digesters identify multiple functional variants in single populations (Ingram et al., 2009b; Jones et al., 2013; Ranciaro et al., 2014): a local soft sweep from multiple mutational origins. Although the regulatory mechanism is not yet fully resolved, it seems that the associated SNPs are loss-of-function mutations (preventing down-regulation) that lead to a gain in enzyme activity (Ingram et al., 2009b). Parallel origin of the adaptive phenotype is facilitated by a large mutational target, leading to a large Θ_*l*_ for adult lactose tolerance.

### Microbes

Some of the most compelling evidence for rapid adaptation comes from microbes, especially from pathogens that evolve drug resistance. We discuss two recombining pathogens in which selective sweeps have been studied: HIV and the Malaria parasite *P. falciparum.*

Drug resistance in HIV typically evolves independently in each patient. Work on selective sweeps in HIV therefore focuses on within-patient populations that start off as susceptible and then acquire one or several resistance mutations. The identity of drug resistance mutations in HIV are well known for all commonly used antiretroviral drugs (Wensing et al., 2015). In the absence of treatment, within-patient HIV populations have large census sizes with around 10^8^-10^9^ active virus-producing cells (Haase, 1994; Coffin and Swanstrom, 2013). Since HIV has a high mutation rate of 10^−7^-10^−5^, depending on mutation type (Abram et al., 2010; Zanini et al., 2016), almost all single nucleotide mutations are present as SGV at any time. With values of Θ_*l*_ ~ 10 to 10^4^, one would expect that adaptation occurs only via soft selective sweeps. However, the scenario may be different for patients under treatment. Although the effect of treatment on the effective population size is unknown, successful treatment can reduce the number of viral particles in the blood by several orders of magnitude.

Paredes et al. (2010) determined whether resistance mutations were already present as SGV in the viral population of 183 patients. Comparing blood samples from before the start of treatment and after treatment had failed, they found that patients with resistance mutations present as SGV were more likely to fail treatment due to resistance evolution (see also Wilson et al., 2016). Another study (Jabara et al., 2011) looked only at one patient, but did very deep and accurate sequencing. The same haplotypes carrying resistance mutations were observed at low frequency in SGV, and at much higher frequency after treatment.

There is also clear evidence for multiple-origin soft sweeps in HIV. The most palpable example comes from patients treated with the drug Efavirenz. The main resistance mutation against this drug is a K103N substitution in the reverse transcriptase gene. The wildtype codon for K (lysine) is AAA. Since codons AAT or AAC both encode N (asparagine), two redundant mutations create the resistant allele. For multiple-origin sweeps, we expect mixtures of AAT and AAC codons in drug resistant populations. Indeed, this is frequently observed (Pennings et al., 2014). However, sometimes only one codon, AAC or AAT, is found. At least some of theses cases are likely hard sweeps. Secondary hardening due to fitness differences (codon bias) among AAC and AAT codons appears unlikely as both versions are observed. A possible explanation is hardening due to linked selection.

Another factor favoring hard sweeps is (temporary) treatment success. Indeed, some patients do not evolve drug resistance at all, or only after many years of treatment (Harrigan et al., 2005; UK Collaborative Group on HIV Drug Resistance, 2005). This suggests a role of *de novo* mutation in resistance evolution. Depending on the patient and the treatment, HIV populations may actually be mutation limited. This is corroborated by a recent analysis of patient derived sequences from the late 1980s until 2013 (Feder et al., 2016). The authors show that the reduction in diversity associated with the fixation of a resistance mutation increases as treatment becomes more effective. This suggests that better treatments lead to harder sweeps, likely because they reduce the population size and move HIV into a mutation-limited regime.

In *Plasmodium falciparum,* unlike in HIV, single drug resistant strains often spread across wide geographic areas. The evolution of drug resistance is thus mostly studied at the level of human populations, rather than the level of parasite populations inside single patients. Patterns of drug resistance in *P. falciparum* roughly come in two forms: When several substitutions are needed to create a resistant allele, observed sweeps are locally hard, but globally soft. When a single substitution or gene duplication event can create a resistant allele, very soft sweeps are observed, both locally and globally.

A striking example of a multiple-origin soft sweep is the evolution of artemisinin resistance due to mutations in the Kelch gene in South-East Asia. Anderson et al. (2017) analysed samples from western Thailand from 2001 (before the spread of artemisinin resistance) through 2014 (when Kelch mutations had nearly fixed). They found 32 different non-synonymous mutations in the Kelch gene, most of which create a resistant phenotype, and estimate a large target size between 87 and 163bp for the resistance allele. However, for the most recent years 2013/14, the authors find that one of the resistance mutations (C580Y) has spread to high frequency (Fig. 1B in Anderson et al., 2017). This has led to partial hardening of the sweep and a lower diversity in samples, likely because of a fitness advantage of C580Y relative to the other resistance mutations. We may see complete hardening of this sweep in future samples. Therefore, as the authors point out, only the observation of a soft sweep in real time allows for the correct assessment of a high risk of resistance evolution. Retrospect analyses after secondary hardening and erroneous classification as a classical hard sweep would miss the essential information.

Another example of a multiple-origin soft sweep comes from resistance evolution to mefloquine and artemisinin due to gene amplification at *pfmdr1* (Nair et al., 2007). The authors find evidence for 5 to 15 independent origins of the gene duplication, based on an analysis of 5' and 3' breakpoints of the amplification events. Since the data was collected two decades after the initial spread of resistance, even more independent origins of this gene amplification may have existed originally.

One example of a locally hard, but globally soft sweep is observed at the *dhfr* gene. Mutations at this gene reduce susceptibility to pyrimethamine. Single, double, and triple mutant haplotypes occur in Africa. In contrast, only one predominant triple mutant haplotype was found in South-East Asia, displaying a classical (local) hard sweep pattern (Nair et al., 2003). The same triple mutant haplotype was found in East and South Africa. Since this haplotype differs from the susceptible or resistant types in Africa, the most parsimonious explanation is that it stems from South-East Asia (Roper et al., 2004). In another study, however, researchers found the same triple mutant on an unrelated genetic background in the West-African archipelago of São Tomé and Principe (Salgueiro et al., 2010), which makes this a globally soft sweep. Another example of a locally hard, but globally soft sweep is found in chloroquine resistance, which is conferred by several non-synonymous mutations in the chloroquine resistance transporter (*crt*) gene (Escalante et al., 2009; Mehlotra et al., 2001).

## 6 Conclusions: Beyond “hard” and “soft”

In a mutation-limited world, recent adaptation leads to hard sweeps and leaves clear and well-understood footprints in genomic diversity. However, existing theory and data indicate that this is not the world we live in. More often than not, adaptation is rapid, resorting to genetic material from various sources, including standing variation, or the recurrent origin of beneficial alleles through mutation or migration.

From theory, we have seen that a single parameter, the population mutation rate Θ = 4*N_e_u*, is most important in separating the rapid world from the mutation limited one. However, this parameter is more complex than it may seem. It depends on the specific beneficial allele and its mutational target size. It also depends on the timing of the adaptive event and the corresponding short-term effective population size. This leads to a large variance for Θ, spanning regions of hard and soft sweeps in most species. Consequently, there is empirical evidence for soft sweeps even in humans, and for hard sweeps even in microbes, despite of huge differences in population size. What is more, the best resolved empirical cases tell us that real adaptive stories go beyond a simplistic hard-or-soft dichotomy.

If adaptation is not mutation limited, we see diversity also among selection footprints. There are classical hard sweeps and two types of “classical” soft sweeps, from a single origin or from multiple origins of the beneficial allele. However, more often then not, real patterns do not fit these model archetypes: For example, there are stacked soft sweeps with secondary hardening, soft sweeps with spatial components, adaptation from pre-adapted variation or from introgression, and there is polygenic adaptation.

There is little reason to argue that either hard sweeps or soft sweeps do not exist. But there is good reason to build on existing concepts and to go on and explore.

## 7 Simulation methods

We simulate a standard Wright-Fisher model of 2*N_e_* haploids. Individual genotypes have m unlinked loci with two alleles each, a wildtype allele *a_i_* and a mutant allele *A_i_*, 1 ≤ *i* ≤ m. Mutation from the wildtype to the mutant occurs at each locus with rate *u*. Although loci are bi-allelic, we distinguish each new mutational origin of a mutant *A_i_* allele from previous mutants at the same locus and record its frequency separately. Let *x_i_* be the total mutant frequency at locus *i*. Prior to *T_S_*, mutant alleles are either neutral or deleterious with selection coefficient *s_d_*. After *T_S_*, each genotype with at least one beneficial allele has selection coefficient *s_b_*. Assuming linkage equilibrium, this results in a marginal fitness of the wildtype of 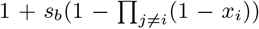. In the life cycle, mutation and selection is followed by independent multinomial sampling at each locus. Prior to *T_S_*, we run the simulation model for 10/*s_d_* generations to approach mutation-selection-drift balance. After *T_S_*, selection turns positive and we follow the trajectories of mutant alleles across all loci. Simulations are stopped once the frequency of the wildtype genotype (with allele *a_i_* at all loci) is < 5%. We record the sweep pattern at the locus with the highest frequency of the beneficial allele at this time, given that this locus has experienced an allele frequency shift of at least 50%.

## 8 Acknowledgments

We thank Graham Coop, Alison Feder, Nandita Garud, Ilse Höllinger, Sebastian Matuszewski, Philipp Messer and an anonymous referee for comments on the manuscript.

